# Insulin preservatives trigger neutrophil extracellular trap formation through SYNE1-mediated nuclear disassembly

**DOI:** 10.64898/2026.06.05.730153

**Authors:** Ulrike Klueh, Ingrida Oendraite, Priscila Silva Cunegundes, Kenneth Wood, Karolis Krinickis, Paul Stemmer, Kaitlin Lowran, Donald L. Kreutzer, Ron Pettis

## Abstract

Neutrophil extracellular traps (NETs) are critical effector molecules in sterile inflammation, yet the molecular mechanisms by which xenobiotic chemical exposures trigger NETosis remain poorly defined. Here, using phenolic preservatives present in all FDA-approved insulin formulations as a discovery platform, we show that these compounds induce NETosis in primary human neutrophils (34.3 ± 5.0% vs. 2.8 ± 0.9% for preservative-free insulin; p < 0.001) via a mechanism distinct from canonical PKC- and calcium-dependent pathways. Data-independent acquisition mass spectrometry (n = 6 donors) reveals that preservatives prompt coordinated dephosphorylation of SYNE1 (nesprin-1) at Ser8724 and Ser8727 (log₂FC = -4.41 and -4.01, respectively; both q-value < 0.0001), disrupting LINC complex-mediated nuclear-cytoskeletal anchoring, through a phosphatase-dependent pathway distinct from canonical PKC- and calcium-dependent NETosis. In a porcine subcutaneous catheter model, preservative-containing formulations drive progressive NET accumulation, neutrophil infiltration, and early fibrotic changes over 7 days, whereas removing preservatives reduces the histological inflammation score by 40% (P < 0.001). These findings establish phenolic preservatives as non-pathogen triggers of NETosis, identify disruption of the SYNE1-LINC complex as the underlying mechanism, and demonstrate that preservative-free formulations lessen device-related inflammation, offering a translatable strategy for safer implantable drug delivery systems.

## INTRODUCTION

Neutrophil extracellular traps (NETs) are increasingly recognized as drivers of sterile inflammation, fibrosis, and implant failure, yet the molecular mechanisms by which nonpathogenic chemical exposures trigger NETosis remain poorly defined. Canonical NETosis pathways involve PKC activation by phorbol esters, calcium flux via ionophores, or pattern recognition of pathogen-associated molecular patterns, all converging on reactive oxygen species production and histone citrullination. Whether xenobiotic small molecules can trigger NETosis through distinct, ROS-independent mechanisms remains to be established. Neutrophils persist at biomaterial implant sites far longer than their canonical role in acute inflammation would predict, as demonstrated by studies showing sustained neutrophil presence and activity at implant interfaces (1–4). Their persistence is associated with ongoing effector functions, including neutrophil extracellular trap (NET) formation, oxidative bursts, protease release, cytokine production, and the formation of prothrombotic scaffolds, potentially contributing to chronic inflammation and tissue damage (2, 3, 5). NETosis is a specialized immune response in neutrophils that traps and kills invading microbes, but it has also been associated with sterile inflammation at implant sites (1). The mechanisms sustaining neutrophil activation in these contexts are increasingly being elucidated, including those related to implant insertion trauma and continuous exposure to biomaterial surface cues, such as topography and stiffness (2, 3). However, their persistence at implant sites and their potential to drive sustained tissue damage through specialized cell death programs, such as NETosis, warrant deeper mechanistic investigation, particularly to inform strategies aimed at modulating neutrophil responses and improving implant outcomes.

Insulin infusion systems provide an experimentally tractable model for investigating preservative-triggered inflammation because they continuously deliver millimolar concentrations of phenolic compounds (m-cresol: 16-30 mM; phenol: 16-29 mM) to subcutaneous tissue. All FDA-approved insulin formulations contain phenolic bacteriostatic preservatives (6), and their pro-inflammatory effects, including neutrophil infiltration, have been characterized in the medical literature. However, some mechanistic details remain under investigation. Phenolic preservatives are added for insulin stability, shelf life, and provide bactericidal properties; they are known to induce cytotoxicity *in vitro* (6, 7), and inflammatory cell infiltration *in vivo* (4–6, 8, 9). However, it remains unclear whether these responses involve NETosis and the signaling mechanisms by which neutrophils detect and respond to preservatives. Critically, one study attributed infusion-site inflammation to insulin protein rather than preservatives (10), highlighting ongoing controversy over the causative agents.

We therefore hypothesized that phenolic preservatives, particularly m-cresol, directly trigger NETosis and thereby sustain chronic inflammation at infusion sites. To test this, we combined functional NETosis assays in primary human neutrophils (9), global phosphoproteomic profiling to define signaling mechanisms, and a clinically relevant porcine catheter model to establish *in vivo* pathological consequences. Here, we show that m-cresol induces NETosis in primary human neutrophils at rates comparable to those of canonical agonists and identify SYNE1 (nesprin-1) dephosphorylation as the central signaling event driving nuclear-cytoskeletal disconnection. *In vivo*, preservative-containing formulations trigger progressive NET deposition and fibrosis, whereas preservative removal attenuates inflammation by 40%. These findings establish phenolic preservatives as a class of xenobiotic NETosis triggers, define a phosphatase-dependent mechanism distinct from pathogen-induced NET formation, and provide strategies for mitigating biomaterial-induced inflammation.

## RESULTS

### Phenolic preservatives trigger distinct neutrophil phosphorylation signatures

Primary human neutrophils were treated with preservative-free insulin [insulin (−)], commercial Humalog® containing preservatives [insulin (+)], or isolated preservatives only (IPP), and analyzed by phosphoproteomic profiling (data-independent acquisition mass spectrometry). Unsupervised (PCA, UMAP) and supervised (PLS-DA) dimensionality reduction revealed no outliers within each group (PCA) and clear separation of treatment groups (PLS-DA and UMAP), demonstrating that preservatives fundamentally reprogram neutrophil phosphorylation landscapes. PLS-DA clearly separated groups in the main comparison between component 1 (16.84%) and 2 (14.02%) and revealed a gradual transition between treatment states when mapping components 2 and 3 (6.19%) (***Fig. 1a***). UMAP separation occurred along the Y-axis, with IPP samples positioned between insulin (+) and insulin (-) samples.

**Figure 1.**
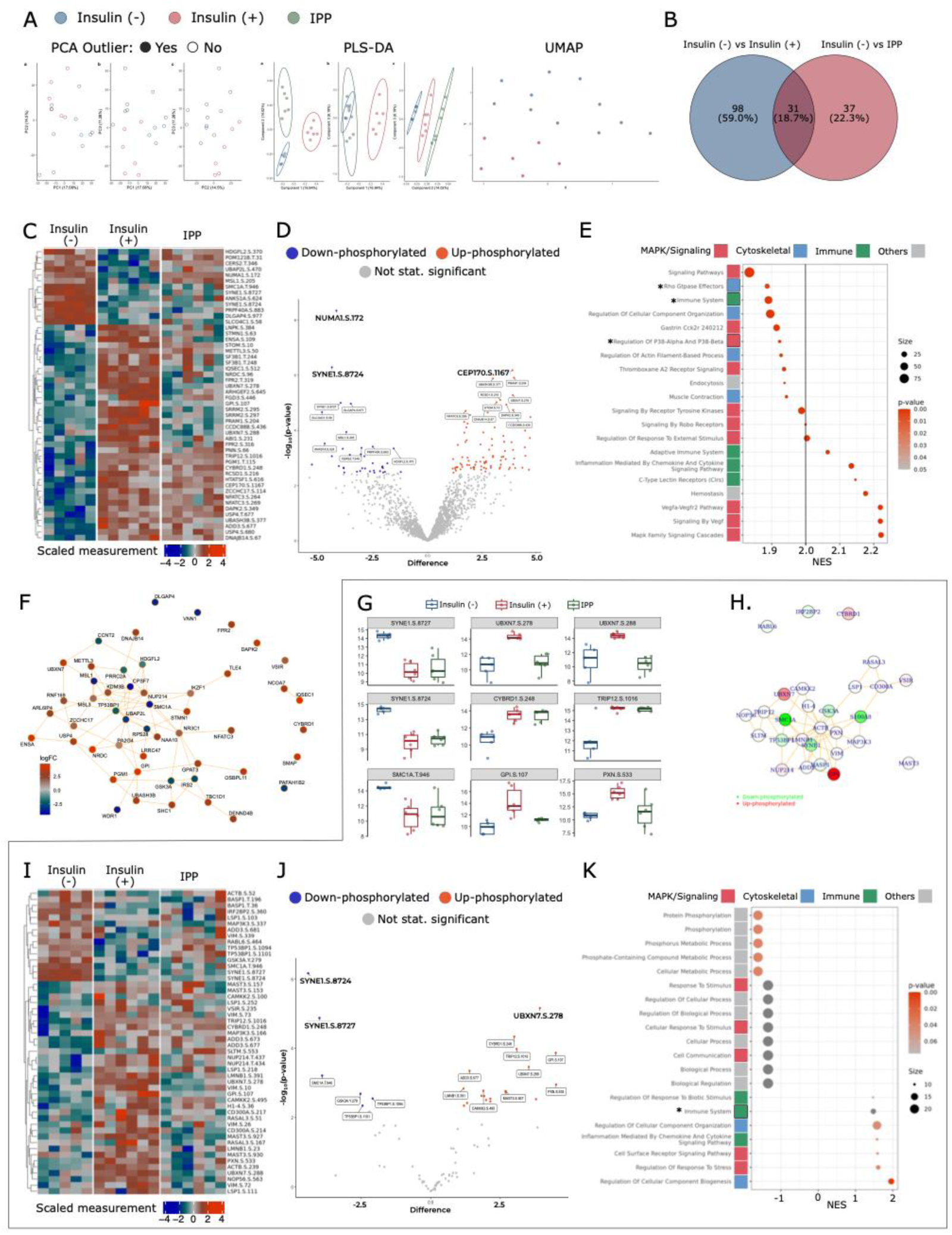
Phenolic preservatives induce distinct phosphoproteomic signatures in human neutrophils. **A**, Dimensionality reduction analysis (PCA, PLS-DA, UMAP) showing dynamics of separation of insulin (+) (red), insulin (−) (blue), and IPP (green) treatment groups. **B**, Venn diagram of significantly regulated phosphosites (|log₂FC| ≥ 0.58, q-value < 0.05) between comparisons. **C**, Heatmap of top 50 phosphosites (insulin (+) vs. insulin (−)). **D**, Volcano plot of differential phosphorylation (insulin (+) vs. insulin (−)). **E**, Pathway enrichment analysis (NES, top 20 pathways). **F**, Protein-protein interaction (PPI) network of top 50 regulated phosphoproteins. **G** to **K** NETosis-related pathway-filtered analysis. **G**, Box plots of representative significantly regulated phosphosites. **H**, PPI network highlighting NETosis-associated modules. **I**, Heatmap of top 50 phosphosites (insulin (+) vs. insulin (−)). **J**, Volcano plot of differential phosphorylation (insulin (+) vs. insulin (−)). **K**, GSEA of ranked phosphoproteins showing enrichment patterns.

Linear modeling with empirical Bayes moderation (|log₂FC| ≥ 1, q-value < 0.05) identified 129 differentially phosphorylated sites across 106 proteins in the insulin (+) versus insulin (−) comparison, comprising 91 upphosphorylated and 38 downphosphorylated sites (***Fig. 1b***). IPP treatment altered 68 phosphosites across 56 proteins (31 up, 37 down). 31 phosphosites (18.7%) were shared between the two contrasts. Hierarchical clustering of the top 50 most significant phosphosites showed opposing patterns between insulin (-) and insulin (+) groups, with IPP displaying an intermediate profile (***Fig. 1c***). Functional classification assigned these phosphosites to five categories: nuclear organization (NUMA1, SMC1A, SYNE1), cytoskeletal regulation (STMN1, ADD3), membrane trafficking (STOM, ARHGEF2), signal transduction (NFATC3, DAPK2), and protein degradation (TRIP12, UBXN7).

Volcano plot analysis identified the largest upphosphorylation changes in cytoskeletal proteins (CEP170 S1167, log₂FC = 3.91; RCSD1 S216, log₂FC = 3.35), and protein degradation machinery (UBXN7 S278, log₂FC = 3.93), with the largest downphosphorylation changes at nuclear envelope proteins (SYNE1 S8727, log₂FC = −4.01; SYNE1 S8724, log₂FC = −4.41; NUMA1 S172, log₂FC = −4.20) and a chromatin remodeling factor (MSL1 S205, log₂FC = −3.75) (***Fig. 1d***). NFATC3 S269 phosphorylation (log₂FC = 1.75) was detected in insulin (+) but not in IPP samples (***Fig. 1j***), suggesting that insulin signaling and preservative toxicity engage distinct regulatory mechanisms.

### Pathway enrichment reveals coordinated activation of NETosis-associated pathways

Gene set enrichment analysis (insulin (+) versus insulin (-) identified enriched pathways spanning MAPK signaling (MAPK Family Signaling Cascades, NES = 2.22, q-value = 0.006; p38α/β Regulation, NES = 1.92), immune activation (Chemokine and Cytokine Signaling, NES = 2.13; C-type Lectin Receptors, NES = 2.15), cytoskeletal reorganization (Rho GTPase Effectors, NES = 1.89; Actin Filament Regulation, NES = 1.93), and hemostasis (NES = 2.18) (***Fig. 1e***). Pathways were color-coded by functional category (red: MAPK/Signaling; green: Immune; blue: Cytoskeletal; gray: Others).

Protein-protein interaction network analysis of the top 50 differentially phosphorylated proteins showed that 40 of 50 nodes formed an interconnected network, with IQSEC1 showing the largest upphosphorylation (logFC 4.74) change and CPSF7 (logFC -4.42) the largest downphosphorylation change (***Fig 1f***). Additional upphosphorylated nodes included NRDC (logFC 4.72), ENSA (logFC 4.48), GPI (logFC 4.50), PGM1 (log FC 3.86), and STMN1 (logFC 3.56). Downphosphorylated nodes included WDR1 (logFC -3.87), SMC1A (logFC -3.91), MSL1 (logFC -3.75), IRS2 (logFC -2.66), and GSK3A (logFC - 2.53).

### Preservatives alter NETosis-relevant phosphorylation independently of insulin

To distinguish preservative-driven from insulin-dependent regulation, we examined six phosphosites selected for NETosis relevance (***Fig. 1g***). NET-relevant phosphorylated proteins, SYNE1 S8727/S8724 (nuclear envelope disruption), SMC1A T946 (chromatin decondensation), CYBRD1 S248 (oxidative metabolism), and TRIP12 S1016 (protein degradation), showed differential regulation between insulin (+) and IPP conditions. UBXN7 S278/S288, GPI S107, and PXN S533 showed differential regulation between insulin (+) and IPP conditions.

Protein-protein interaction network analysis (50 nodes, 83 edges, PPI enrichment p = 0.027) showed upregulation of GPI, UBXN7, LMNB1, NUP214, CD300A, and VSIR, and downregulation of S100A8, SYNE1, and TP53BP1 (***Fig. 1h***). ACTB occupied a central position with minimal phosphorylation change.

Quantitative heatmap analysis showed high baseline phosphorylation of SYNE1 S8724 and S8727 in insulin (-) samples and reduced phosphorylation in both IPP and insulin (+) samples (***Fig. 1i***). SMC1A T946 showed a similar pattern conditions. LMNB1 S331 and NUP214 showed intermediate phosphorylation in IPP samples and higher phosphorylation in insulin (+) samples.

### SYNE1 dephosphorylation is the most statistically significant phosphorylation change

Volcano plot analysis showed that SYNE1 S8724 (log₂FC = −4.41, q-value < 0.0004) and S8727 (log₂FC = −4.01, q-value < 0.002) were the most statistically significant dephosphorylation events in the insulin (+) versus insulin (-) comparison (***Fig. 1j***). Protein-protein interaction network analysis showed that SYNE1 connects to ACTB within the cytoskeletal scaffold (***Fig. 1h***).

Pathway enrichment analysis (insulin (+) versus (insulin (-) identified two downregulated phosphorylation-associated gene sets: Protein Phosphorylation, NES = -1.60, p =0.04 and Phosphorylation, NES = -1.61, p = 0.04 (***Fig. 1k***). SYNE1 S8724 and S8727 dephosphorylation was greater in insulin (+) than in IPP samples (***Fig. 1g***)

### Preservatives induce robust NET formation in primary human neutrophils

To validate that the identified phosphorylation signature corresponds to functional NETosis, we performed quantitative imaging and flow cytometry of primary human neutrophils treated with preservative-containing and preservative-free formulations. Live-cell imaging (four independent experiments, quadruplicate wells) demonstrated progressive extracellular DNA release over 12 h in insulin (+), IPP, and m-cresol (5 μM) conditions, quantified by Cytotox Green fluorescence (***Fig. 2a-c***). Insulin (-) and untreated controls showed minimal cytotoxicity. Dual-parameter analysis (nuclear intensity versus Cytotox Green) revealed a characteristic NETosis profile, progressive increases in both nuclear expansion and membrane permeabilization, distinct from apoptotic or necrotic cell death (***Fig. 2c***).

**Figure 2.**
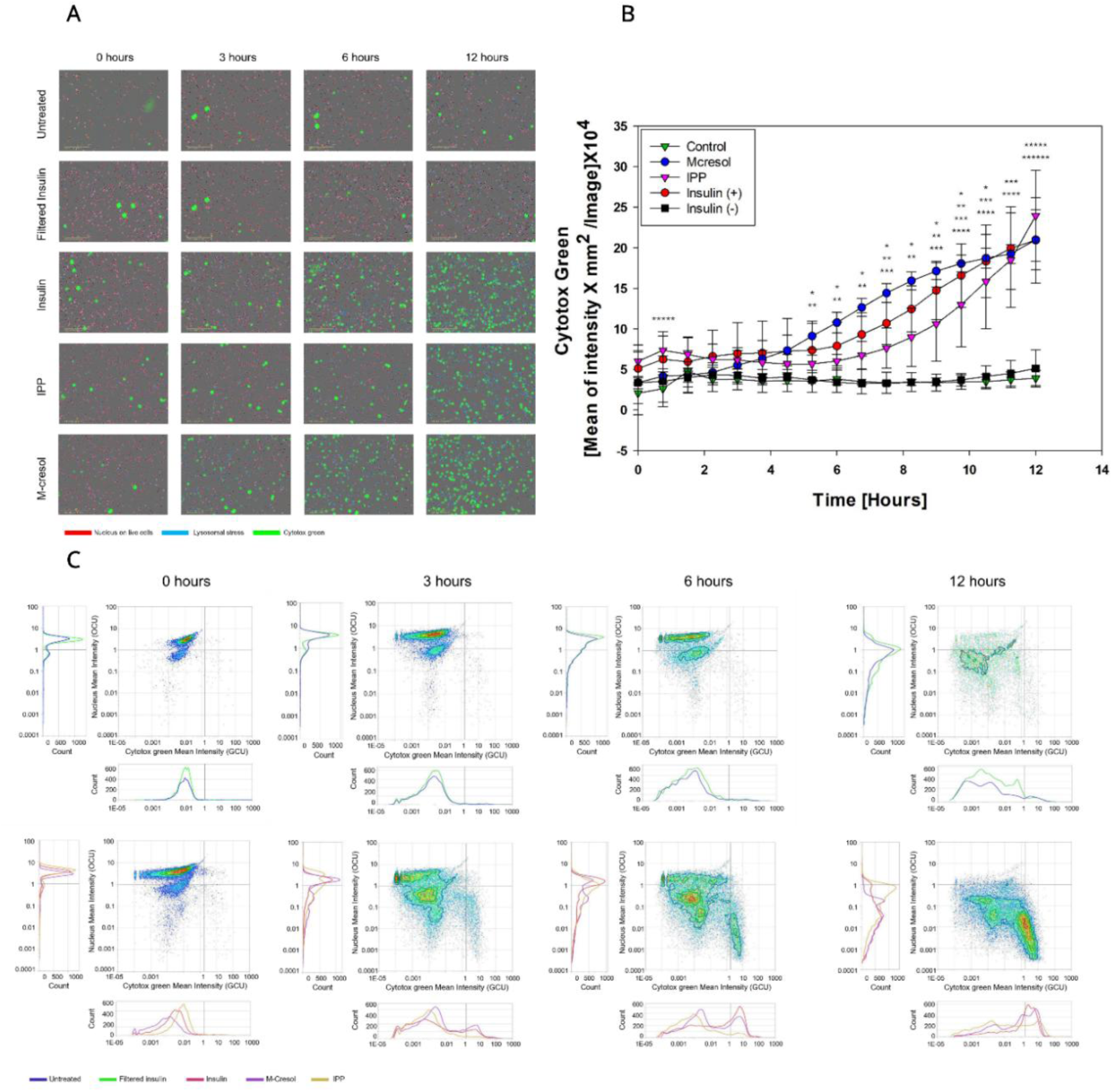
Incucyte- time-lapse monitoring was performed every 45 minutes for 12 hours on untreated and treated human neutrophils. Human neutrophil treatment included insulin (+), IPP, M-cresol, and insulin (-). **A)** Representative images were taken at 0, 3, 6, and 12 hours. Nucleus are identified in blue and DNA from disrupted cells in green. **B)** Quantification of the cytotoxic effects of treatments was performed, expressing the mean intensity of cytotox green, including the standard deviation. This was calculated from the micrometer square of nine images per well, using five independent experiments run in triplicate. Statistical significance was determined by One-Way ANOVA for each treatment, treating replicate values as independent data points and using a 95% confidence interval (p>0.05). Mcresol vs. Control **\***, Mcresol vs. Filtered Insulin **\*\***, Insulin vs. Filtered Insulin **\*\*\***, Insulin vs. Control **\*\*\*\***, IPP vs. Control **\*\*\*\*\*,** IPP vs. Filtered Insulin **\*\*\*\*\*\***. **C)** Representative dot plots and histograms were created for each axis, showing the population distribution between the X-axis (cytotox green mean intensity) and the Y-axis (nucleus mean intensity) at time points 0, 3, 6, and 12 hours. Colored lines on the histograms represent the different treatments: the first dot plot line is Untreated (blue) and insulin (-) (green). Second dot plot’s line Insulin (+) (red), M-cresol (purple), and IPP (yellow).

Flow cytometry confirmed that preservative-containing formulations induced NETosis at rates comparable to established agonists: insulin (+) 34.3 ± 5.0%, IPP 35.0 ± 4.3%, m-cresol 35.6 ± 4.2%, versus ionophore A23187 38.2 ± 4.8% (all p < 0.001 versus controls) (***Fig. 3a-c***). NETosis was defined by dual criteria: (1) nuclear decondensation (>2-fold increase in area: insulin (+) 156 ± 12 μm², IPP 149 ± 11 μm², m-cresol 162 ± 14 μm² versus untreated 45 ± 6 μm²; all p < 0.0001), and (2) MPO-positive extracellular DNA (Manders’ coefficient: insulin (+) 0.78 ± 0.06, IPP 0.76 ± 0.05, m-cresol 0.81 ± 0.07 versus ionophore 0.85 ± 0.05; all p < 0.001 versus controls) (***Fig. 3b,c-f***). Preservative-treated neutrophils also secreted high levels of IL-8 (plateauing at 1,000 pg ml^-1^ by 6 h), whereas insulin (-) showed no significant IL-8 production (***Fig. 3h***), confirming functional neutrophil activation.

**Figure 3.**
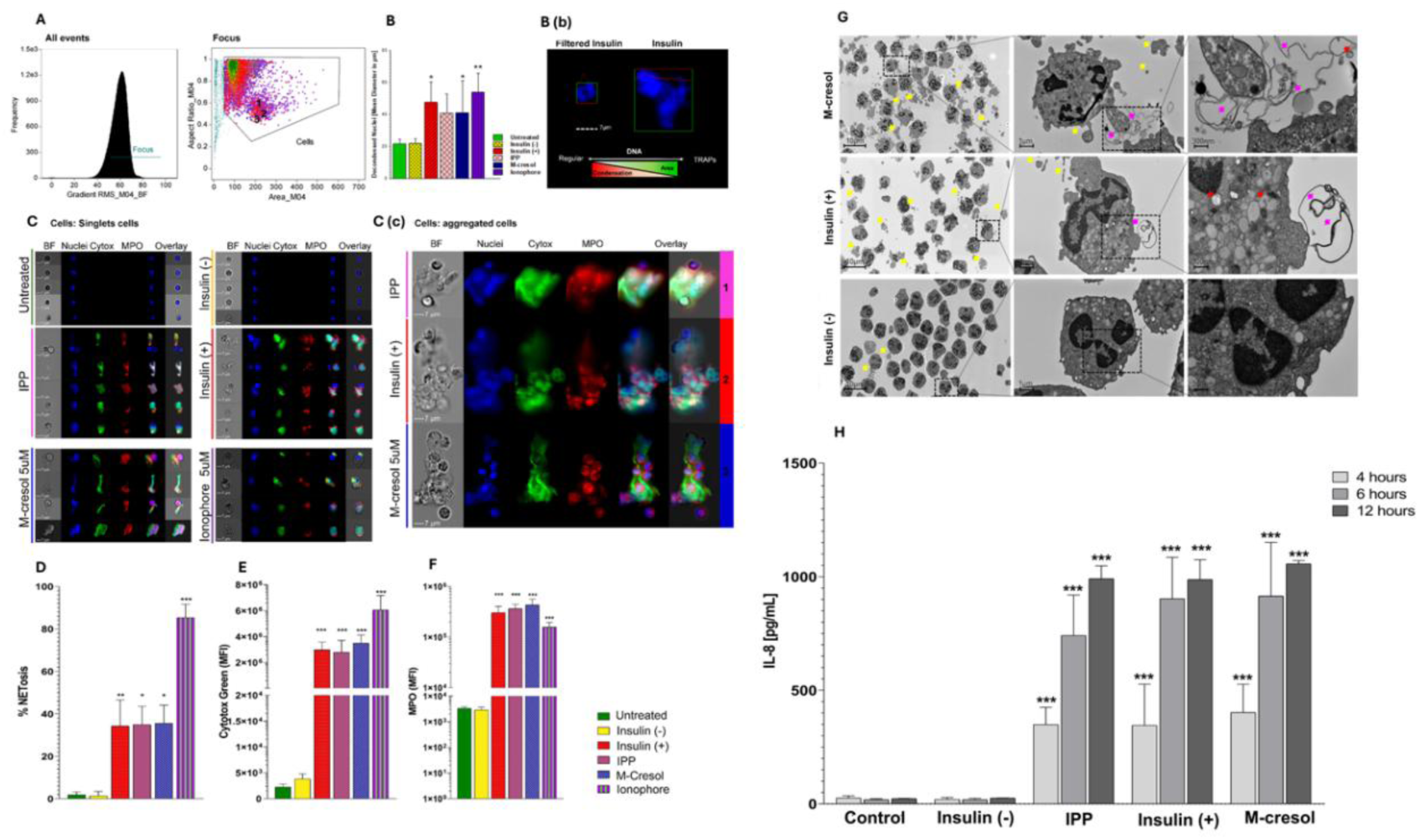
M-Cresol Induces Neutrophil Extracellular Trap Formation and Functional Activation. Human neutrophils were analyzed to assess NET formation, nuclear morphological changes, and functional activation in response to preservatives in insulin formulations. Imaging flow cytometry (panels A-F) captured 10,000 events per sample at 60× magnification and analyzed using IDEAS 6.2 software. Transmission electron microscopy (panel G) and cytokine secretion assays (panel H) confirmed preservative-induced NET formation and neutrophil activation. **(A)** Gating strategy for singlet and aggregated cell populations. Events from all samples were gated using the default M04 mask to generate histogram-gradient RMS-M04-BF, identifying focused events on the brightfield channel. Dot plots display area (x-axis) versus aspect ratio/circularity (y-axis) of brightfield (M04 mask) to discriminate singlet from aggregated cells. Colored dots represent different treatments: green, untreated neutrophils; yellow, filtered insulin; pink, insulin phenolic preservative (IPP); red, insulin; blue, m-cresol; purple, ionophore (positive control). **(B)** Quantification of nuclear morphology changes within singlet cell populations. Bar graph shows mean nuclear diameter (µm) across treatments. **(b)** Representative images of singlet cells from filtered insulin and insulin treatments, highlighting nuclear morphological differences. Scale bar, 7 µm. Schematic illustrates DNA condensation and area differences between a regular nucleus and NETosis. **(C)** Representative singlet events from each treatment showing brightfield (BF), nuclei (Hoechst 5 µM), cytotox green, myeloperoxidase (MPO-NL637), and merged channels (1, Nucleus+Cytotox+MPO; 2, all channel. **(c)** Representative aggregated cell events from IPP, insulin, and m-cresol treatments, with corresponding regions indicated in the dot plot (panel A). **(D)** Percentage of NETosis calculated as the proportion of double-positive events for cytotox green and MPO. **(E)** Mean fluorescence intensity (MFI) of cytotox green across treatments. **(F)** Mean fluorescence intensity (MFI) of MPO across treatments. **(G)** Transmission electron microscopy of human neutrophils (2 × 10⁶ cells/mL) treated with m-cresol (5 µM), insulin (+) (Humalog, 1:6 dilution), or insulin (−) (phenolic preservative-depleted, 1:6 dilution) for 2 h at 37°C, 5% CO₂. Images shown at three magnifications per treatment (left to right: 10 µm, 1 µm, 300 nm). Yellow arrows, extracellular vesicles and cellular debris; pink arrows, extracellular DNA traps (present only in m-cresol and insulin (+) samples); red arrows, autophagic vacuoles containing DNA fragments. Representative images from three independent experiments examining ≥100 cells per condition. **(H)** IL-8 secretion by preservative-treated neutrophils. Preservative-treated neutrophils secreted high levels of IL-8, plateauing at 1,000 pg ml⁻¹ by 6 h, whereas insulin (−) showed no significant IL-8 production, confirming functional neutrophil activation.

Critically, preservative-free insulin [insulin (-)] exhibited minimal NETosis (2.8 ± 0.9%, nuclear area 48 ± 7 μm², MPO colocalization 0.15 ± 0.04), statistically indistinguishable from untreated controls (2.1 ± 0.8%, p = 0.45), definitively establishing that preservatives, not insulin protein, drive NET formation. Transmission electron microscopy (representative images from three independent experiments) confirmed NETosis hallmarks in m-cresol and insulin (+) samples: chromatin decondensation, nuclear membrane disruption, and extracellular fibrillar DNA networks (***Fig. 3g***). Autophagosome-like structures containing DNA fragments were observed in m-cresol- and insulin (+)-treated cells but were absent in insulin (-)-treated cells.

### Preservatives drive progressive NET deposition and fibrosis in vivo

To establish clinical relevance, we evaluated tissue responses to continuous m-cresol buffer infusion (3.15 mg ml⁻¹) (HE (+)) in a porcine catheter model over 3, 5, and 7 days (***Fig. 4a-j***). The sample size (n=4) is consistent with prior published studies (9, 11–13) and this porcine study was designated as a proof-of-concept pilot. Quantitative histopathology (five-point scoring) revealed minimal inflammation at day 3 (median score 1.5), significant escalation by day 5 (3.75, p = 0.05 versus day 3), and sustained chronic inflammation through day 7 (3.75, p = 0.01 versus day 3) (***Fig. 4k,l***). Immunofluorescence staining for citrullinated histone H3 (H3Cit) and myeloperoxidase (MPO) demonstrated progressive NET accumulation paralleling the inflammatory trajectory: isolated NET foci at day 3, substantial deposition by day 5, and extensive high-intensity NET signals by day 7 (***Fig. 4m-o***).

**Figure 4.**
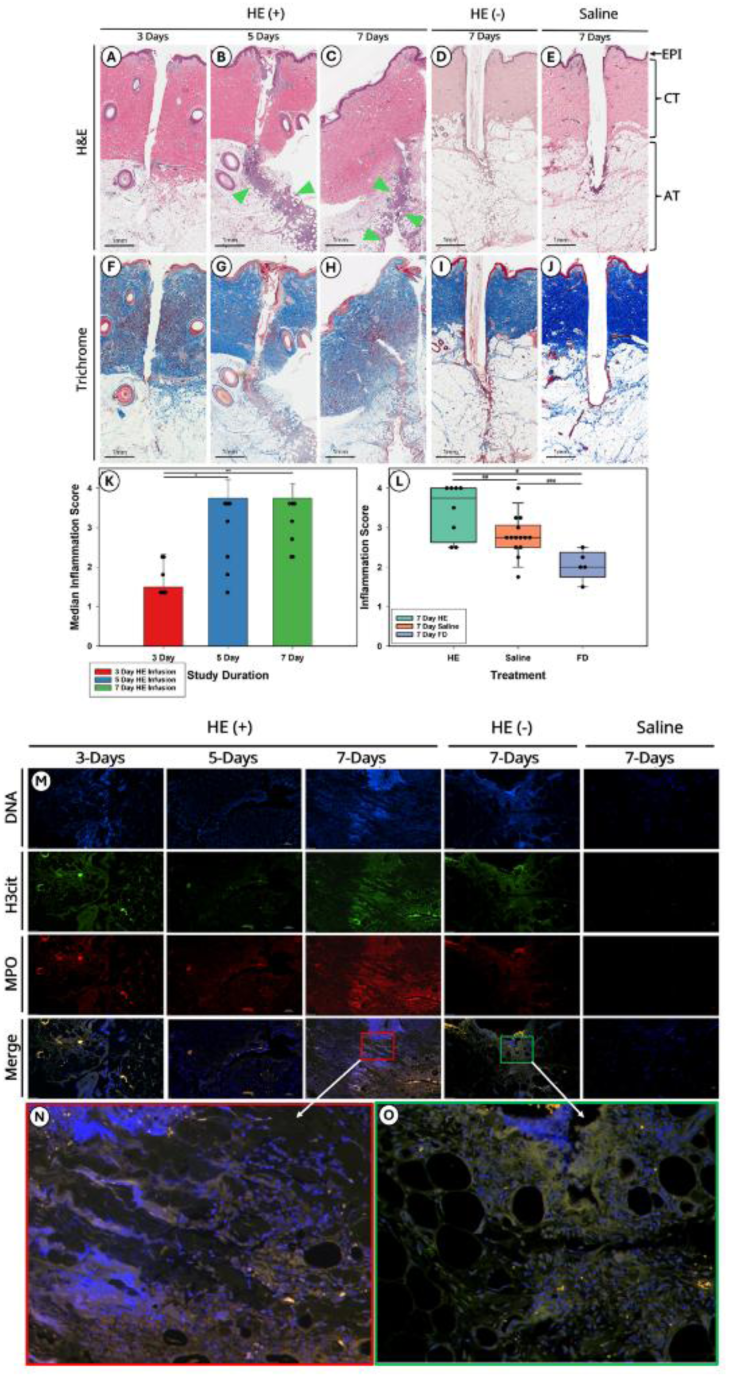
Time-Dependent Tissue Inflammation and NET Formation at Insulin Preservative Infusion Sites. Swine subcutaneous tissue sections showing time-dependent inflammatory responses to m-cresol containing buffer (HE (+)) infusion. Tissue architecture shows epidermis (EPI), connective tissue (CT), and adipose tissue (AT). Histological analysis by hematoxylin and eosin (H&E) and Masson’s trichrome staining; immunofluorescence detection of NET components by confocal microscopy**. (A-C)** H&E-stained sections of m-cresol (Humalog excipient (HE (+)) infusion sites at 3 days (A), 5 days **(B)**, and 7 days **(C)**. Green arrows denote inflammatory infiltration pathways from the infusion site through the tissue layer. **(D)** H&E-stained section of 7-day buffer (HE (-)) infusion site, showing reduced inflammatory response compared to saline and m-cresol. **(E)** H&E-stained section of 7-day saline infusion site (negative control). **(F-J)** Masson’s trichrome-stained sections corresponding to panels A–E, showing collagen deposition (blue) and fibrotic tissue formation around implant sites at 3 days **(F)**, 5 days **(G)**, 7 days IPP **(H)**, 7 days filtered **(I)**, and 7 days saline **(J)**. **(K)** Quantification of median inflammatory scores over time for IPP infusion. Inflammatory scores increased progressively from 3 to 7 days. Data represent median values from histological scoring of tissue sections (n = 4 animals per time point). P < 0.05, P< 0.03, 3-day versus 5-day and 7-day IPP, respectively (one-way ANOVA with post hoc tests). **(L)** Comparison of 7-day tissue inflammation scores between m-cresol (Humalog excipient), saline control, and buffer infusion. HE (+) infusion induced significantly greater inflammation than both saline and HE (-) controls. Data represent median ± interquartile range (n = 4 animals per group). #P < 0.05, ##P < 0.01, ###P < 0.001 (one-way ANOVA with 95% confidence interval and post hoc comparisons as indicated). **(M)** Wide-field fluorescence microscopy of NET components at infusion sites over time. Tissue sections from panels **(A – E)** were immunostained for NET markers: DNA (DAPI, blue), citrullinated histone H3 (H3Cit-FITC, green), and myeloperoxidase (MPO-NL637, red). Images show progressive NET accumulation at the infusion release sites from 3 to 7 days. Merged images show overlay of all fluorescence channels. **(N,O)** High-magnification comparison of NET formation at 7-day infusion sites. Merged fluorescence images (boxed regions from panel M) comparing m-cresol **(N)** and buffer **(O)** infusions, demonstrating reduced NET formation with preservative removal. Scale bars: 1 mm. Representative images from at least three independent experiments with 4 animals per group.

To determine whether inflammation was preservative-specific, we compared the m-cresol-free buffer (HE (-)) and saline at day 7. m-Cresol infusion induced the highest inflammation score (3.44), followed by saline (2.79) and preservative-free buffer (2.05) (one-way ANOVA, p < 0.001) (***Fig. 4l***). Removal of m-cresol reduced inflammation by 40.4% (p < 0.001) and virtually eliminated NET formation (***Fig. 4l***), establishing proof of concept for preservative removal as a therapeutic strategy. Saline-treated controls showed negligible NET deposition, confirming that preservatives, not mechanical catheter trauma alone, drive sustained NETosis.

Trichrome staining revealed progressive peri-catheter collagen deposition in m-cresol-treated (HE (+)) sites: minimal deposition at day 3, discrete collagen fibers by day 5, and organized fibrous capsule formation by day 7 (Fig. 4f–h). Preservative-free (HE (-)) and saline controls exhibited minimal collagen deposition through 7 days (Fig. 4l,j), suggesting that preservative-induced NETosis is linked to early fibrotic tissue remodeling.

## DISCUSSION

We identify SYNE1 dephosphorylation as a non-canonical mechanism of xenobiotic-induced NETosis, and establish m-cresol, a phenolic preservative present in all FDA-approved insulin formulations, as the trigger driving nuclear-cytoskeletal disconnection (***Fig 1***). In primary human neutrophils, m-cresol induces NETosis at rates comparable to those of canonical agonists (35% versus 2.8% for preservative-free insulin, p < 0.001) (***Fig 2, 3***), and in a clinically relevant porcine model, preservative-containing formulations drove progressive NET accumulation, sustained inflammation, and early fibrosis (Fig 4a-j), with preservative removal reducing inflammation by 40% (p < 0.001), providing proof of concept for preservative-free formulation strategies in implantable drug delivery (***Fig 4l***). The intermediate inflammation score observed with saline (2.79) suggests a procedural or catheter-related baseline inflammatory component (***Fig 4l***), underscoring that the comparison between m-cresol and preservative-free buffer specifically isolates the preservative-driven contribution. These findings have direct clinical relevance given the widespread use of phenolic preservatives in insulin formulations.

Mechanistically, m-cresol-induced NETosis diverges from canonical pathways. Whereas PMA and calcium ionophores activate PKC- and calcium-dependent signaling (14–16), respectively, m-cresol triggers coordinated dephosphorylation of SYNE1 S8724/S8727 (***Fig. 1j***), the most statistically significant change identified. SYNE1 (nesprin-1) anchors the nuclear envelope to the actin cytoskeleton through LINC complexes; its dephosphorylation disrupts this anchoring, enabling nuclear disassembly and chromatin extrusion. This phosphatase-dependent mechanism establishes phenolic preservatives as a functionally distinct trigger class.

### SYNE1 dephosphorylation as a regulatory node in nuclear disassembly

Nuclear envelope breakdown is rate-limiting in NETosis (17–20), but the upstream signals governing LINC complex disassembly remain poorly defined (21–23). SYNE1 forms the cytoplasmic component of LINC complexes, which span the nuclear envelope to connect chromatin-associated SUN proteins with the actin cytoskeleton (24, 25). While SYNE1 is critical for nuclear positioning and mechanotransduction, phosphorylation of SYNE1 is not established as a mechanism maintaining these processes during interphase (26, 27); our finding that m-cresol triggers SYNE1 dephosphorylation at multiple residues (38 downregulated nuclear/chromatin proteins identified) suggests coordinated phosphatase activation as a regulatory mechanism for nuclear-cytoskeletal disconnection. This contrasts with infection-induced NETosis, in which neutrophil elastase and myeloperoxidase drive nuclear envelope breakdown through proteolytic degradation (28). The identification of this phosphatase-dependent pathway provides a new entry point for investigating how chemical irritants interface with neutrophil nuclear architecture.

Kinase substrate enrichment analysis revealed coordinated dephosphorylation within the Protein Phosphorylation pathway in insulin (+) samples (NES = -1.60, p = 0.04), consistent with phosphatase activation contributing to nuclear destabilization, including at SYNE1. This explains the synergistic effect observed in commercial formulations and suggests that other agents activating phosphatases may similarly enhance preservative-induced NETosis. The specific phosphatase(s) responsible remain to be identified, but candidates include protein phosphatase 1 (PP1) and protein phosphatase 2A (PP2A), which regulate nuclear envelope dynamics during mitosis (29–31).

### Phenolic preservatives expand the repertoire of NETosis triggers

Phenolic preservatives expand the repertoire of NETosis triggers beyond PAMPs, DAMPs, and immune complexes to include small-molecule xenobiotics. At millimolar concentrations (16-30 mM in commercial formulations), m-cresol induces NETosis with kinetics and magnitude comparable to those of canonical agonists, yet operates through a distinct phosphatase-dependent mechanism. This has implications for understanding sterile inflammation in contexts where synthetic chemicals accumulate in tissue, including drug-eluting devices, topical formulations, and environmental exposures.

The molecular sensor(s) mediating m-cresol recognition remain unidentified, but candidates include the aryl hydrocarbon receptor (AhR), which binds aromatic compounds and regulates immune responses (32–34), and transient receptor potential (TRP) channels, which sense chemical irritants and trigger calcium influx (35). The hyperphosphorylation (inactivation) of NFATC3 Ser2009 in insulin-containing formulations, but not with preservatives alone, further distinguishes this mechanism from canonical calcium-NFAT-dependent NETosis. Canonical pathogen-induced NETosis involves calcium influx, calcineurin activation, and NFAT dephosphorylation to drive transcriptional programs including inflammatory cytokine production. In contrast, preservative-induced NETosis proceeds through direct nuclear envelope disruption (SYNE1 dephosphorylation) without NFAT activation, indicating a transcription-independent, post-translational mechanism. This may explain the relatively lower inflammatory cytokine production observed with chemical NETosis compared to pathogen-triggered NETosis, as only IL-8 was increased in the measured cytokines (***Fig 3h***). Preservative-induced IL-8 secretion (1,000 pg ml⁻¹ by 6 h) likely contributes to sustained neutrophil recruitment, creating a positive feedback loop that amplifies inflammation (**Fig. 3h**).

### NETosis-driven biomaterial inflammation: implications beyond insulin delivery

These findings have relevance beyond insulin delivery. Many implantable devices, including drug-eluting stents, intrathecal pumps, and extended-release depots, expose tissues to sustained concentrations of preservatives and excipients, yet their capacity to trigger NETosis has not been systematically evaluated. Our temporal analysis, revealing progressive NET accumulation from day 3 to day 7 *in vivo* (Fig. 4m), suggests that chronic, low-level NETosis may drive the transition from acute inflammation to fibrotic encapsulation observed in foreign body responses. This is consistent with emerging literature implicating NETs in fibrosis across multiple organ systems, where NET-associated proteases (elastase, cathepsin G, matrix metalloproteinases) degrade extracellular matrix and NET-DNA scaffolds serve as templates for collagen deposition (36–39).

The observation that preservative-free insulin formulations exhibited minimal NETosis (2.8%) and reduced inflammation (40% reduction, p < 0.001) establishes proof of concept that targeting NETosis can improve biomaterial biocompatibility. Potential strategies include: (1) reformulation to minimize or eliminate phenolic preservatives; (2) development of in-line preservative-removal devices for continuous infusion systems; (3) adjunctive pharmacological inhibition of NETosis pathways (for example, PAD4 inhibitors, which prevent histone citrullination required for chromatin decondensation (40, 41)). Beyond device applications, our findings suggest that phenolic compounds present in topical medications and encountered in industrial settings may contribute to contact dermatitis and, potentially, chronic skin inflammation through NETosis- associated mechanisms.

These findings may have particular relevance beyond insulin delivery. This may be especially relevant for the growing group of subcutaneously administered GLP-1 receptor agonists (e.g., semaglutide, liraglutide), which also contain phenolic preservatives like phenol and m-cresol. Injection site reactions are a known adverse effect of these drugs, but the role of phenolic preservatives in causing these reactions is not fully understood (7, 42). Investigating whether NETosis-related pathways contribute to these reactions requires further research.

### Reconciling preservatives versus insulin as inflammatory triggers

Our findings establish preservatives as the primary NETosis trigger, resolving previous uncertainty about the relative contributions of insulin protein, preservatives, and mechanical trauma (9, 11, 42–45). Although one study attributed infusion-site inflammation to insulin rather than preservatives (10), that analysis relied on bulk inflammatory cell counts rather than assessing NETosis-specific endpoints. The equivalence of NETosis rates induced by m-cresol alone (35.6%), IPP extract (35.0%), and commercial insulin (34.3%), contrasted with minimal NET formation by preservative-free insulin (2.8%), provides definitive evidence for preservative-driven NETosis. This does not exclude contributions from insulin aggregates or catheter materials to overall inflammation; indeed, our previous work demonstrated that device insertion induces transient NET formation (9).

### Limitations and Future Directions

Several questions remain. As noted above, the molecular sensor(s) responsible for m-cresol recognition, including AhR and TRP channel candidates, remain to be identified. Separately, the specific phosphatase(s) that drive SYNE1 dephosphorylation remain to be identified. Our phosphoproteomics study, like many others, faces the challenge that the functional roles of the vast majority of identified phosphosites remain unknown. Even the largest human phosphoproteome dataset, encompassing over 119,000 phosphosites, leaves most unannotated (43). This makes it difficult to determine whether observed changes, such as those in SYNE1 phosphorylation, represent regulatory events central to the m-cresol response or incidental bystander modifications. Therefore, future work should employ targeted functional approaches, such as phosphomimetic and phosphodead mutagenesis or CRISPR-based editing of specific SYNE1 phosphosites, to determine which events are biologically consequential and identify the upstream phosphatases responsible for SYNE1 dephosphorylation.

Additionally, we isolated neutrophils from healthy donors, and we used non-diabetic swine; whether diabetes-associated metabolic perturbations (hyperglycemia, neutrophil priming) modulate preservative-induced NETosis warrants investigation, as neutrophils from individuals with diabetes exhibit altered functional responses (9, 46–51). The swine model used a small sample size (n = 4 per group), consistent with prior infusion pump studies but insufficient to detect subtle genotype-phenotype interactions (9, 11–13). Finally, while we focused on m-cresol as the most cytotoxic preservative, phenol and other excipients (for example, polysorbates, benzyl alcohol) warrant systematic evaluation for NETotic potential.

### Concluding Perspective

We establish m-cresol as a small-molecule NETosis inducer that operates through SYNE1-mediated nuclear-cytoskeletal disconnection, a mechanism distinct from pathogen-triggered NET formation. *In vivo*, preservative-containing formulations drive progressive NET deposition and fibrosis, whereas preservative removal attenuates inflammation by 40%, establishing proof of concept for therapeutic intervention. Beyond insulin delivery, these findings have implications for a range of implantable devices that expose tissues to preservatives and excipients. The SYNE1-LINC complex axis represents a tractable target for mitigating biomaterial-induced inflammation, and the signaling pathways defined here provide a framework for investigating how chemical irritants interface with innate immune effector programs.

## MATERIALS AND METHODS

### Study Design and Sex as biological variable

This study employed a multimodal experimental design combining *in vivo* porcine catheter models with *in vitro* human neutrophil studies to investigate tissue reactions and cellular responses to m-cresol, a phenolic preservative commonly found in insulin formulations. Our swine study examined male and female swine, and the findings are reported for both sexes.

### Animal Studies Animal and Housing

Healthy, non-diabetic Yucatan swine (Sinclair; aged 4-6 months, 25-35 kg) were housed under a 12:12-hour light/dark cycle at 20-22°C, with ad libitum water and twice-daily feeding. Sample size was determined using a power calculation based on published tissue inflammation endpoints in swine catheter studies, where effect sizes typically range from 50–65% between device types, with standard deviations of 15–30% of the mean (Cohen’s d = 1.67–4.33) (10, 13, 45). Assuming 6 devices per animal, an intraclass correlation coefficient of 0.2 for within-animal clustering, 80% power, and α = 0.05, we determined that 4 animals per group would provide adequate power to detect moderate to large differences in tissue response.

### Catheter Implantation

Six MiniMed Quick-set infusion sets with 6 mm 90° catheters (Medtronic) were implanted per animal using randomized placement to minimize location bias. Histopathological evaluations were performed by blinded investigators.

### Formulation

Test formulations (Humalog® excipient containing no insulin, referred to as HE (+)) contained m-cresol (3.15 mg/mL), dibasic sodium phosphate (1.88 mg/mL), zinc (19.7 µg/mL), glycerin (16 mg/mL), in sterile water, pH adjusted to 7.4 ± 0.1. Control formulations included 0.9% saline and excipient without m-cresol, designated as HE (-). All formulations were prepared under sterile conditions, stored at 4°C, and used within 48 hours after confirming pH stability and sterility.

### Infusion and Tissue Collection

Formulations were continuously infused at 22.5 µL/h for 3, 5, or 7 days using programmable infusion pumps. Animals were monitored daily for signs of distress, infection, or catheter displacement. Eight-millimeter punch biopsies were collected 24 hours post-catheter removal, cleaned with antiseptic solution, and immediately fixed either in Formalin or Zinc Fixative (BD Pharmingen) for histopathological analysis.

### Histopathological and Immunofluorescence

#### Tissue Processing and Staining

Biopsies were fixed for 24 hours, processed through graded alcohols, and embedded in paraffin. Sections (5 µm) were stained with hematoxylin and eosin (H&E) or Masson Trichrome (Epredia).

### NET Detection by Immunofluorescence

Sections were deparaffinized, rehydrated, and subjected to antigen retrieval in citrate buffer (pH 6.0) at 95°C for 20 minutes. After blocking with 5% normal goat serum in PBS for 1 hour, sections were incubated overnight at 4°C with primary antibodies: anti-histone H3 (Abcam ab3594, 1:500) and anti-elastase (Abcam ab68672, 1:200). FITC-conjugated secondary antibodies (Abcam ab7086, Cambridge, UK, 1:1000) were applied for 2 hours at room temperature. Nuclei were counterstained with DAPI (Electron Microscopy Sciences Fluoro-Gel II). For MPO detection, anti-MPO antibody (R&D Systems AF3667, 1:300) followed by anti-goat IgG-NL637 (R&D Systems NL002, 1:500) was used.

### Tissue Reactions Scoring

H&E-stained sections from epidermis (EPI), connective tissue (CT), and adipose tissue (AT) were scored using a standardized 5-point scale (0 = no reaction, 1 = minimal, 2 = mild, 3 = moderate, 4 = severe) for inflammation, fibrosis, and cellular infiltration. Two independent, blinded observers analyzed ≥ 5 high-power fields (400×) per section. Inter-observer reliability (ICC > 0.85) was confirmed.

### Human Neutrophil Studies

#### Neutrophil Isolation

Venous blood (4 ml) was collected from healthy adult donors (n=6) in K2EDTA Vacutainer tubes (Greiner Bio-One) and processed within 2 hours using the EasySep Direct Human Neutrophil Isolation kit (StemCell Technologies, Cambridge, MA). Purity (>95%) was confirmed by cytospin at 600 rpm for 3 minutes (Cyto-Tek®, Sakura). Viability (>90%) was assessed using a Bio-Rad TC20 automated cell counter with trypan blue exclusion. Neutrophils were resuspended at 1×10⁶ cells/mL in phenol-free RPMI-1640 medium supplemented with 10% FBS.

### Formulation Preparation for *In Vitro* Studies

Phenolic compounds were removed from Humalog® (Eli Lilly & Co., Indianapolis, IN, USA) using Zeba spin desalting columns (7K MWCO, Thermo Fisher Scientific, Waltham, MA, USA) pre-equilibrated with three DPBS washes. Humalog® (700 µL) was centrifuged at 1,000 × g for 2 minutes at room temperature. Eluted insulin free of phenolic compounds [insulin (−)], unprocessed Humalog [insulin (+)], and wash fractions containing insulin phenolic preservatives (IPP; concentration confirmed by Prussian Blue assay) were collected. M-cresol solution (3.15 mg/mL in PBS) was filter-sterilized (0.22 µm). Treatments were diluted 1:6 in culture medium, corresponding to a final m-cresol concentration of 5 µM in samples containing m-cresol.

### Real-time NET Visualization

Primary human neutrophils (1×10⁵ cells/well) were seeded in clear-bottom black 96-well plates (Corning) in phenol-free RPMI medium with 10% FBS and pre-loaded with Cytotox Green (Sartorius, Göttingen, Germany, Cat# 4633, 250 nM) for extracellular DNA detection and NucSpot® Live 650 (Fremont, CA, USA, Cat# 40082) for nuclear staining. After treatment with insulin (+), insulin (−), IPP, or m-cresol (1:6 dilution, 5 µM final m-cresol), plates were immediately transferred to the IncuCyte SX5 live-cell (Sartorius, Göttingen, Germany) analysis system, and cells were imaged every 45 minutes for 12 hours. NET formation was quantified as mean fluorescence intensity per µm² per image (9 images per well, ≥ 500 cells/condition) across four independent experiments performed in quadruplicate.

### Image Flow Cytometry Analysis

Neutrophils (2×10⁶ cells per 200 µL) were seeded in U-bottom 96-well polypropylene plates (Cellstar/Greiner) and treated with 250 nM Cytotox Green and incubated for 2 hours at 37°C and 5% CO₂. Untreated cells served as negative controls; cells treated with 5 µM calcium ionophore A23187 (Sigma-Aldrich, St. Louis, Mo, USA) served as positive controls. Cells were fixed with 4% paraformaldehyde in 0.1% BSA buffer for 20 minutes at 4°C, washed three times with DPBS, and blocked with 1% BSA in DPBS for 30 minutes. After overnight incubation at 4°C with goat anti-MPO (R&D Systems AF3667, 1:200) on an orbital shaker, cells were incubated with Donkey anti-goat IgG NorthernLights™ NL637 (R&D Systems, Cat# NL002, 1:250) and Hoechst 33342 (Invitrogen, Carlsbad, CA, USA, 5 µM) for 2 hours at room temperature. Samples were acquired on an ImageStream® X Mark II imaging flow cytometer (Amnis, Seattle, WA, USA) using INSPIRE acquisition software, and data were analyzed with IDEAS software. NET-positive cells, defined as cells with extracellular DNA co-localizing with MPO, were identified using automated algorithms (minimum of 10,000 neutrophils per condition).

### Cytokine Analysis

Supernatants collected at 4, 6, and 12 hours were analyzed for IL-10, IL-12p70, IL-1β, IL-33, IL-4, IL-6, IL-8, MCP-1, and TNF-α using electrochemiluminescence detection (MSD U-Plex human kit) on the QuickPlex SQ120, following the manufacturer’s instructions.

### Electron Microscopy

Neutrophils (2 × 10⁶ cells/mL) in RPMI with 10% FBS were treated with PBS (control), insulin (+), insulin (−), IPP, or m-cresol (1:6 dilution) for 2 hours at 37°C with 5% CO₂. Cells were centrifuged (1,100 rpm, 5 min) and immediately fixed with double-concentrated fixative (4% paraformaldehyde and 4% glutaraldehyde in 0.2 M cacodylate buffer, pH 7.4) at 1:1 ratio for 2 h at 4°C. Pellets were washed three times with 0.1 M cacodylate buffer, post-fixed with 2% OsO₄ and 3% K₃Fe(CN)₆ in 0.1 M cacodylate buffer (1 h on ice), and washed five times with double-distilled water. Samples were sequentially stained with 1% thiocarbohydrazide (20 min, room temperature), 2% OsO₄ in water (40 min), 1% uranyl acetate (4°C, overnight), and Walton’s lead solution (30 min, 60°C). Samples were dehydrated through graded ethanol (50%, 70%, 90%, 100%) with increasing resin concentrations, embedded in EMBed-812/DER 736 resin with activator, and polymerized at 60°C for 48 hours. Ultrathin sections (70-80 nm) were cut with a Leica ARTROS 3D ultramicrotome with a diamond knife and imaged on a Zeiss Gemini 300 scanning electron microscope equipped with a backscatter detector. Representative images were acquired at standardized magnifications (10,000× and 50,000×) to visualize cellular ultrastructure, nuclear morphology, extracellular DNA release, and changes in subcellular organelles. A minimum of 100 cells per condition were examined across three independent experiments by blinded investigators to assess morphological features consistent with NETosis.

### Phosphoproteomics

Global phosphoproteomic profiling of primary human neutrophils (n=6 donors) treated with three formulations: preservative-free insulin [insulin (-)], commercial Humalog containing preservatives [insulin (+)], and phenolic preservatives used in Humalog (IPP).

### Sample Preparation

Neutrophils (10⁶ cells/well) from six healthy donors were treated with insulin (−), insulin (+), or IPP at 1:6 dilution in RPMI-1640 with 10% FBS for 30 minutes at 37°C with 5% CO₂. After three washes with 14 mL ice-cold PBS and centrifugation (1,000 rpm, 5 minutes, 4°C), cell pellets were flash-frozen in liquid nitrogen and stored at -80°C. Pellets were solubilized in 200 µL of 2.5% lithium dodecyl sulfate at 95 °C for 5 min and clarified using spin columns (Pierce, Cat# 89868). Protein concentration was determined by BCA assay (Pierce, Cat# 23235). Samples were buffered with 20 mM tetraethylammonium bicarbonate, reduced with 5 mM dithiothreitol (37 °C, 30 min), alkylated with 15 mM iodoacetamide (room temperature, 30 min, dark), and quenched with 5 mM dithiothreitol. Proteins were precipitated with methanol, resuspended in 20 mM Tris (pH 8.0), 10 mM CaCl₂, and 10% acetonitrile, and digested with trypsin (Sigma-Aldrich, Cat# EMS0007; 1:50 enzyme-to-protein ratio) at 47 °C for 2 h followed by overnight incubation at 37 °C.

### Phosphopeptide Enrichment and Mass Spectrometry

Phosphopeptides were enriched using Fe-NTA magnetic beads (Cube Biotech, Cat# 31525-FE). Dried peptides were reconstituted in 80% acetonitrile with 0.1% trifluoroacetic acid, incubated with 20 µL pre-washed beads, and processed using a KingFisher robotic platform. Beads were washed three times with 0.1% formic acid in 80% acetonitrile, and phosphopeptides were eluted with 2% ammonium hydroxide in 50% acetonitrile. Eluates were neutralized to pH 3 with 10% formic acid, dried, and reconstituted in 0.1% formic acid.

LC-MS/MS was performed using a Thermo Scientific Vanquish Neo HPLC equipped with an Acclaim PepMap 100 trap column (100 µm × 2 cm, C18, 5 µm, 100Å) and an Ion Optics Aurora analytical column (75 µm × 25 cm) maintained at 45°C. A 120-minute gradient from 1% to 42% solution B (80% acetonitrile, 0.1% formic acid) was applied at 300 nL/min. Data-independent acquisition was conducted on a Thermo Scientific Orbitrap Eclipse mass spectrometer. MS1 spectra were acquired at 120,000 resolution (350-1200 m/z), AGC target of 3 × 10⁶, and maximum injection time of 50 ms. DIA windows covered 400-1000 m/z with 8 m/z isolation windows at 15,000 resolution.

### Data Processing

Data were processed using Spectronaut 19.3 (Biognosys) with default normalization and local regression normalization against the UniProt human reference proteome (UP000005640, downloaded March 30, 2021). Trypsin digestion with up to two missed cleavages was specified. Fixed modification: carbamidomethylation of cysteine. Variable modifications: phosphorylation (S, T, Y), methionine oxidation, and N-terminal acetylation. The false discovery rate was set at 1% for both precursor and peptide levels. Quality control metrics included peptide retention time stability (<2% CV), mass accuracy (<5 ppm), and protein identification reproducibility (>80% overlap) across technical replicates. Phosphopeptide intensities were log-transformed and retained only if data were available for at least 50% of samples in each condition. Missing values were imputed in a site- and condition-specific manner only when ≥50% of data were present for a given condition. Principal Component Analysis identified outlier samples (>3 standard deviations from the mean in the first three principal components); due to the limited sample size, outliers were retained. Data were normalized by median centering without scaling using PhosR v1.12.0 (52).

### Bioinformatic data analysis

#### Phosphoproteomics

Differential phosphorylation was assessed using limma v3.58.1 (53). A linear model (∼ 0 + Substance) was fitted by least-squares with lmFit, followed by empirical Bayes moderation with eBayes. P-values were adjusted using the Benjamini-Hochberg procedure to correct for false discovery rate (FDR) and a q-value < 0.05 was considered significant. Differentially expression criteria: |log₂(fold-change)| ≥ 0.58 (1.5-fold) and q-value < 0.05. Pathway enrichment analysis was performed using clusterProfiler v4.10.1 (54) with Bader Lab database (downloaded November 4, 2024) for pathways with 10-500 genes. Signalome analysis was performed using STRINGdb v2.14.0 (55) with the STRING database API (v12.0, species 9606). The top 50 most significant phosphoproteins were mapped to protein-protein interaction networks, with nodes colored by log fold change. NETosis-associated residue analysis was performed using Gene Set Enrichment Analysis with fgsea v1.34.2. Residues were ranked by signed log p-values, and human pathways with keywords NEUTROPHIL, DEGRANULATION, GRANULE, CHROMATIN, ROS, and PAD4 were filtered for NETosis specific filtering.

### General Statistics

Normality was assessed using the Shapiro-Wilk test (n < 50) or Kolmogorov-Smirnov test (n ≥ 50). Variance homogeneity was evaluated using Levene’s or Brown-Forsythe test. For normally distributed data with equal variances, one-way ANOVA with Tukey’s post hoc test (multiple comparisons) or unpaired t-test (two groups) was used. For normally distributed data with unequal variances, Welch’s ANOVA with Games-Howell post hoc test or Welch’s t-test was used. For non-normally distributed data, Kruskal-Wallis test with Dunn’s post hoc test (multiple comparisons) or Mann-Whitney U test (two groups) was used. Tissue observational analysis used Pearson product-moment correlation for reliability. Power analysis was conducted for medium effect sizes (Cohen’s d = 0.5) with 80% power at α = 0.05. Post hoc power analyses were performed for all primary endpoints. Data are presented as mean ± SEM unless otherwise specified. Significance was set at p < 0.05; exact p-values are reported when p > 0.001. Two-tailed tests were used unless specifically justified. Statistical analyses were performed using GraphPad Prism 9.0, SigmaPlot 15, and R version 4.3.0.

### Study approval

Animal studies were conducted in accordance with Wayne State University Institutional Animal Care and Use Committee (IACUC). Human studies received approval from the Wayne State University Institutional Review Board (IRB); all participants provided written informed consent.

## Data Availability

Raw data supporting the conclusions are available from the corresponding author upon reasonable request. Proteomics data have been deposited in the ProteomeXchange Consortium via the PRIDE partner repository under accession number (Pending).

## Author contributions

**U.K.** Conceptualization, funding acquisition, overall coordination of the study, writing of the first draft, and final manuscript.

**I.O. and K. K.**: Performing phosphoproteomics bioinformatics analysis and interpretation, along with writing the phosphoproteomics results.

**K.W.**: Experimental execution of swine studies, imaging, and statistical analyses.

**P.M.S.** and **K.L.:** Phosphopeptide enrichment and mass spectrometry data acquisition; writing of the corresponding Materials and Methods sections.

**P.S.C.:** Finalization of manuscript figures and critical review of the manuscript

**R.J.P.:** Contributed to the conceptual design of the swine studies.

**D.L.K.:** Contributed to the histopathologic analysis of the swine studies and critical review of the manuscript.

## Acknowledgments

We acknowledge the assistance of the Wayne State University Proteomics Core, which is supported through NIH grants P30ES036084, P30CA022453, and S10OD030484. The authors wish to thank Dr. Jean G. de Souza for his contributions to the experimental execution of IncuCyte, flow cytometry, and transmission electron microscopy assays, and for his participation in drafting portions of the Materials and Methods and Results sections related to these experiments. His early contributions to the experimental framework of this study are appreciated. Carol Atkinson, Executive Director of Insulin for Life USA, provided the infusion set supplies. The authors thank Ms. Li Mao for histopathology assistance.

## Artificial intelligence (AI)-assisted tools (like LLMs)

AI-assisted literature search and synthesis platforms, including Open Evidence and SciSpace, were used to support the identification and review of relevant publications. All cited sources were independently verified by the authors. AI-assisted tools, including Claude (Anthropic, version Sonnet 4.6) and Gemini (Google, version 3.1 Pro), were used during manuscript preparation for language editing, sentence refinement, and writing clarity. All scientific content, data interpretation, and conclusions were generated and verified by the authors. The authors take full responsibility for the integrity and accuracy of the work. All AI-assisted edits were reviewed, evaluated, and approved by the authors prior to inclusion.

## REFERENCES

1. Barbeck M, Jung O. Granulocytes and their Involvement in the Foreign Body Response to Biomaterials and Tissue Repair. In Vivo. 2026;40(1):1–16.

2. Clancy DM, et al. Extracellular Neutrophil Proteases Are Efficient Regulators of IL-1, IL-33, and IL-36 Cytokine Activity but Poor Effectors of Microbial Killing. Cell Rep. 2018;22(11):2937–2950.

3. Dömer D, et al. Neutrophil Extracellular Traps Activate Proinflammatory Functions of Human Neutrophils. Front Immunol. 2021;12:636954.

4. Dorai VK, et al. A biomaterial implant model demonstrates that immature neutrophils drive immunopathology following acute injury. Biomaterials. 2026;328:123907.

5. Hidalgo A, et al. Neutrophil extracellular traps: from physiology to pathology. Cardiovasc Res. 2022;118(13):2737–2753.

6. Kesserwan S, et al. Advancing continuous subcutaneous insulin infusion in vivo: New insights into tissue challenges. J Biomed Mater Res A. 2021;109(7):1065–1079.

7. Weber C, et al. Phenolic excipients of insulin formulations induce cell death, pro-inflammatory signaling and MCP-1 release. Toxicol Rep. 2015;2:194–202.

8. Abaricia JO, et al. Control of innate immune response by biomaterial surface topography, energy, and stiffness. Acta Biomater. 2021;133:58–73.

9. Wood KA, et al. Injury to tissue caused by device penetration of the skin triggers formation of extracellular traps. Commun Biol. 2025;8(1):1669.

10. Swinney MR, et al. Insulin, Not the Preservative m-cresol, Instigates Loss of Infusion Site Patency Over Extended Durations of CSII in Diabetic Swine. J Pharm Sci. 2021;110(3):1418–1426.

11. Cunegundes PS, et al. Phenolic Preservatives Are Not the Sole Cause of Eosinophilic Infiltration at Infusion Pump Sites. Diabetes Technol Ther. 2025;27(8):577–586.

12. Hauzenberger JR, et al. Systematic in vivo evaluation of the time-dependent inflammatory response to steel and Teflon insulin infusion catheters. Sci Rep. 2018;8(1):1132.

13. Hauzenberger JR, et al. Detailed Analysis of Insulin Absorption Variability and the Tissue Response to Continuous Subcutaneous Insulin Infusion Catheter Implantation in Swine. Diabetes Technol Ther. 2017;19(11):641–650.

14. Altman A, Mally MI, Isakov N. Phorbol ester synergizes with Ca2+ ionophore in activation of protein kinase C (PKC)alpha and PKC beta isoenzymes in human T cells and in induction of related cellular functions. Immunology. 1992;76(3):465–471.

15. Takai Y, et al. Role of protein kinase C in transmembrane signaling. J Cell Biochem. 1985;29(2):143–155.

16. Wolf M, et al. A model for intracellular translocation of protein kinase C involving synergism between Ca2+ and phorbol esters. Nature. 1985;317(6037):546–549.

17. Li Y, et al. Nuclear envelope rupture and NET formation is driven by PKCα-mediated lamin B disassembly. EMBO Rep. 2020;21(8):e48779.

18. Neubert E, et al. The power from within - understanding the driving forces of neutrophil extracellular trap formation. J Cell Sci. 2020;133(5):jcs241075.

19. Reis LR, et al. Citrullination of actin-ligand and nuclear structural proteins, cytoskeleton reorganization and protein redistribution across cellular fractions are early events in ionomycin-induced NETosis. Redox Biol. 2023;64:102784.

20. Thiam HR, et al. NETosis proceeds by cytoskeleton and endomembrane disassembly and PAD4-mediated chromatin decondensation and nuclear envelope rupture. Proc Natl Acad Sci. 2020;117(13):7326–7337.

21. Cain NE, et al. Conserved SUN-KASH Interfaces Mediate LINC Complex-Dependent Nuclear Movement and Positioning. Curr Biol CB. 2018;28(19):3086–3097.e4.

22. Cruz VE, Esra Demircioglu F, Schwartz TU. Structural Analysis of Different LINC Complexes Reveals Distinct Binding Modes. J Mol Biol. 2020;432(23):6028–6041.

23. King MC. Dynamic regulation of LINC complex composition and function across tissues and contexts. FEBS Lett. 2023;597(22):2823–2832.

24. Jahed Z, et al. Molecular models of LINC complex assembly at the nuclear envelope. J Cell Sci. 2021;134(12):jcs258194.

25. Sosa BA, Kutay U, Schwartz TU. Structural insights into LINC complexes. Curr Opin Struct Biol. 2013;23(2):285–291.

26. Espigat-Georger A, et al. Nuclear alignment in myotubes requires centrosome proteins recruited by nesprin-1. J Cell Sci. 2016;129(22):4227–4237.

27. Graham DM, Burridge K. Mechanotransduction and nuclear function. Curr Opin Cell Biol. 2016;40:98–105.

28. Tabrizi ZA, et al. Multi-facets of neutrophil extracellular trap in infectious diseases: Moving beyond immunity. Microb Pathog. 2021;158:105066.

29. Holder J, Poser E, Barr FA. Getting out of mitosis: spatial and temporal control of mitotic exit and cytokinesis by PP 1 and PP 2A. FEBS Lett. 2019;593(20):2908–2924.

30. Li J, et al. Mechanisms of PP2A-Ankle2 dependent nuclear reassembly after mitosis. eLife. 2025;13:RP104233.

31. Mehsen H, et al. PP2A-B55 promotes nuclear envelope reformation after mitosis in Drosophila. J Cell Biol. 2018;217(12):4106–4123.

32. Larigot L, et al. Aryl Hydrocarbon Receptor and Its Diverse Ligands and Functions: An Exposome Receptor. Annu Rev Pharmacol Toxicol. 2022;62:383–404.

33. Larsson M, et al. Identification of potential aryl hydrocarbon receptor ligands by virtual screening of industrial chemicals. Environ Sci Pollut Res Int. 2018;25(3):2436–2449.

34. Polonio CM, et al. The aryl hydrocarbon receptor: a rehabilitated target for therapeutic immune modulation. Nat Rev Drug Discov. 2025;24(8):610–630.

35. Mager ML, et al. TRPA1-dependent and -independent activation by commonly used preservatives. Front Pharmacol. 2023;14:1248558.

36. Chrysanthopoulou A, et al. Neutrophil extracellular traps promote differentiation and function of fibroblasts. J Pathol. 2014;233(3):294–307.

37. Sorvillo N, et al. Extracellular DNA NET-Works With Dire Consequences for Health. Circ Res. 2019;125(4):470–488.

38. Wu X, Yang Y. Neutrophil extracellular traps (NETs) and fibrotic diseases. Int Immunopharmacol. 2024;133:112085.

39. Zheng Y, et al. The Roles of Neutrophil Cells in the Pathogenesis of Fibrosis: Mechanisms for Disease Progression. FASEB J Off Publ Fed Am Soc Exp Biol. 2025;39(17):e70992.

40. Dakin LA, et al. Inhibiting peptidylarginine deiminases (PAD1-4) by targeting a Ca2+ dependent allosteric binding site. Nat Commun. 2025;16(1):4579.

41. Gajendran C, et al. Alleviation of arthritis through prevention of neutrophil extracellular traps by an orally available inhibitor of protein arginine deiminase 4. Sci Rep. 2023;13(1):3189.

42. Kalus A, et al. Evaluation of Insulin Pump Infusion Sites in Type 1 Diabetes: The DERMIS Study. Diabetes Care. 2023;46(9):1626–1632.

43. Kesserwan S, et al. Inflammation at Site of Insulin Infusion Diminishes Glycemic Control. J Pharm Sci. 2022;111(7):1952–1961.

44. Kesserwan S, et al. A pharmacological approach assessing the role of mast cells in insulin infusion site inflammation. Drug Deliv Transl Res. 2022;12(7):1711–1718.

45. Kastner JR, et al. In Vivo Study of the Inflammatory Tissue Response Surrounding a Novel Extended-Wear Kink-Resistant Insulin Infusion Set Prototype Compared With a Commercial Control Over Two Weeks of Wear Time. J Diabetes Sci Technol. 2023;17(6):1563–1572.

46. Carestia A, et al. NETosis before and after Hyperglycemic Control in Type 2 Diabetes Mellitus Patients. PLOS ONE. 2016;11(12):e0168647.

47. Farhan A, et al. Spontaneous NETosis in diabetes: A role of hyperglycemia mediated ROS and autophagy. Front Med. 2023;10:1076690.

48. Jin L, et al. Neutrophil extracellular traps (NETs)-mediated killing of carbapenem-resistant hypervirulent *Klebsiella pneumoniae* (CR-hvKP) are impaired in patients with diabetes mellitus. Virulence. 2020;11(1):1122–1130.

49. Joshi MB, et al. Glucose induces metabolic reprogramming in neutrophils during type 2 diabetes to form constitutive extracellular traps and decreased responsiveness to lipopolysaccharides. Biochim Biophys Acta BBA - Mol Basis Dis. 2020;1866(12):165940.

50. Joshi MB, et al. Elevated homocysteine levels in type 2 diabetes induce constitutive neutrophil extracellular traps. Sci Rep. 2016;6(1):36362.

51. Soongsathitanon J, Umsa-Ard W, Thongboonkerd V. Proteomic analysis of peripheral blood polymorphonuclear cells (PBMCs) reveals alteration of neutrophil extracellular trap (NET) components in uncontrolled diabetes. Mol Cell Biochem. 2019;461(1–2):1–14.

52. Kim HJ, et al. PhosR enables processing and functional analysis of phosphoproteomic data. Cell Rep. 2021;34(8):108771.

53. Ritchie ME, et al. limma powers differential expression analyses for RNA-sequencing and microarray studies. Nucleic Acids Res. 2015;43(7):e47–e47.

54. Yu G, et al. clusterProfiler: an R package for comparing biological themes among gene clusters. Omics J Integr Biol. 2012;16(5):284–287.

55. Szklarczyk D, et al. The STRING database in 2023: protein–protein association networks and functional enrichment analyses for any sequenced genome of interest. Nucleic Acids Res. 2023;51(D1):D638–D646.

